# Circular inference in bistable perception

**DOI:** 10.1101/521195

**Authors:** Pantelis Leptourgos, Charles-Edouard Notredame, Marion Eck, Renaud Jardri, Sophie Denève

**Affiliations:** Laboratoire de Neurosciences Cognitives & Computationnelles, ENS, INSERM U960, PSL Research University, 75005, Paris, France; Univ Lille, CNRS UMR-9193, SCALab, PsyCHIC team & CHU Lille, Fontan Hospital, CURE Platform, F-59037 Lille, France

**Keywords:** bistability, Necker cube, Bayesian inference, circular inference

## Abstract

When facing fully ambiguous images, the brain cannot commit to a single percept and instead switches between mutually exclusive interpretations every few seconds, a phenomenon known as bistable perception. Despite years of research, there is still no consensus on whether bistability, and perception in general, is driven primarily by bottom-up or top-down mechanisms. Here, we adopted a Bayesian approach in an effort to reconcile these two theories. Fifty-five healthy participants were exposed to an adaptation of the Necker cube paradigm, in which we manipulated sensory evidence (by shadowing the cube) and prior knowledge (e.g., by varying instructions about what participants should expect to see). We found that manipulations of both sensory evidence and priors significantly affected the way participants perceived the Necker cube. However, we observed an interaction between the effect of the cue and the effect of the instructions, a finding incompatible with Bayes-optimal integration. In contrast, the data were well predicted by a circular inference model. In this model, ambiguous sensory evidence is systematically biased in the direction of current expectations, ultimately resulting in a bistable percept.

## Introduction

Perception can be defined as the process of combining available information to create valid and useful interpretations of the world. Although our phenomenological experience makes us think that perceptual decisions are trivial, the truth might be very different. An interesting example is visual perception of depth. When we see an object, our brain must reconstruct its 3D shape from a 2D retinal image; in other words, the brain must solve an inference problem [1]. Unfortunately, such problems are ill-posed, as in most cases the 2D retinal projection is compatible with many different 3D objects [2]. To cope with perceptual uncertainty, the brain must combine ambiguous information received by peripheral sensors (e.g., disparity cues and movement cues) with pre-existing information (either hard-wired or learned) concerning properties of the environment or the potential cost of a wrong decision [3, 4]. Such combinations can be expressed through Bayes’ theorem, in which prior probability distributions and sensory likelihoods are multiplied, resulting in a posterior probability distribution over possible solutions to the perceptual problem. Most of the time, only a single dominant (most probable) interpretation will emerge from these constraints.

However, when the level of ambiguity is too high, finding a single interpretation is not possible. Strikingly, ambiguous figures compatible with more than one plausible interpretation [5, 6] lead to *bistable* (or more generally *multistable)* perception [7]. When facing those figures, the perceptual system is unable to commit to a single stable interpretation and instead oscillates between mutually exclusive interpretations every few seconds. A famous figure known to induce bistability is the *Necker cube* (NC) ([5]; **Figure 1a**), in which a 2D collection of lines is automatically interpreted as a 3D cube, which is either “seen from above” (SFA interpretation) or “seen from below” (SFB interpretation). Interestingly, the NC is an asymmetrical stimulus, meaning that it generates an implicit preference for the SFA interpretation (i.e., the general preference of humans to interpret things as if they were below the level of their eyes) [3, 8].

**Figure 1:**
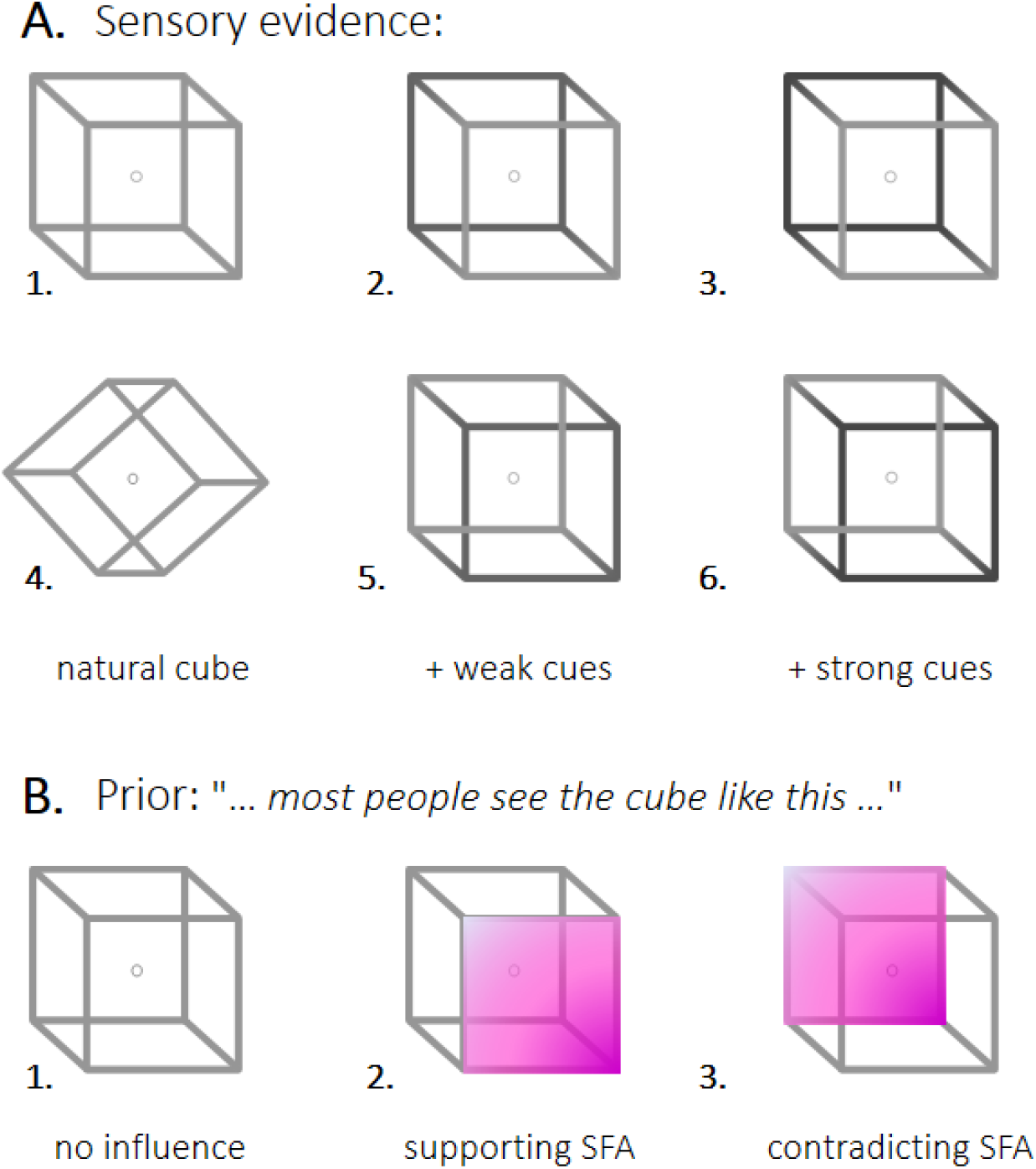
Stimuli and instructions. **(a)** Different Necker cubes were used to induce bistable perception, in which the 2D figure is perceived as a 3D cube with either the left or the right side closer to the observer. Even in the case of the completely ambiguous stimulus **(1)**, people have an implicit preference to interpret the cube as seen from above (SFA interpretation), which was interpreted an implicit prior. This prior can be annihilated by tilting the stimulus **(4)**. Sensory evidence was manipulated by adding visual cues in the form of contrasts **(2-3,5-6)**. The contrast could be strong **(3,6)** or weak **(2,5)** and could support **(2,3)** or contradict **(5,6)** the implicit prior. **(b)** A further manipulation of the prior was achieved by giving correct or wrong information to the participants about which interpretation was generally stronger (explicit prior). Instructions could support or contradict the implicit prior. An additional control group received no particular instructions. Crucially, to avoid additional priming effects, all groups received the same visual instructions (including the stimulus and the 2 possible interpretations), and the differences were only verbal. Note that the color used in the present figure has only been added for illustration purposes; during the experiment, participants were presented with full cubes.

While the concept of perception as inference under uncertainty offers a principled way to explain the efficiency of perceptual systems and certain perceptual illusions, it can account for bistable perception less directly. Indeed, if the brain uses explicit representations of uncertainty, e.g., a probability distribution instead of a point estimate [9–12], ambiguous stimuli should be recognized as such and not generate a unique, persistent representation. We note that bistable perception is far from unique in that case. Although many studies have reported that the brain is able to reach Bayes-optimal decisions [13–16], there are numerous tasks in which human behavior deviates significantly from that of a Bayesian observer [17–20].

Deviations from Bayesian optimality could be the consequence of highly non-linear and state-dependent interactions between feedback and feedforward streams of information in brain circuits [21]. Some of these effects can be quantified by the *circular inference* framework [22]. According to this framework, hierarchical processing in the brain is analogous to the propagation of probabilistic messages (beliefs) in a hierarchical model of the world [23]. The combination of feedforward and feedback inputs is equivalent to the product of prior and likelihood in Bayes’ theorem. However, because neural circuits are highly recurrent, sensory evidence and prior information can easily reverberate and be artificially amplified through feedforward/feedback loops in the brain, resulting in the corruption of sensory evidence by prior information and *vice versa.* Such reverberation can be avoided if excitation (E) and inhibition (I) are perfectly balanced in cortical circuits [22], a well-known property of the healthy brain [24, 25].

Recently, our team hypothesized a link between E/I imbalance in schizophrenia and the occurrence of psychotic symptoms (hallucinations and delusions). This hypothesis was recently reinforced by experimental evidence in a probabilistic reasoning task [26]. Interestingly, we also detected a certain amount of circularity in healthy participants, particularly the corruption of sensory evidence by prior information. If circular inference is a more general mechanism than initially predicted, an interesting question arises: is it possible to find evidence of circularity [27] in the perceptual behavior of healthy subjects in the absence of any psychotic experience? Here, we propose that bistability could be an example of percepts induced by such circularity.

To investigate this theory, we induced bistability in healthy participants using the NC. We asked how different pieces of information, including (a) pre-existing priors (i.e., the SFA preference), (b) newly acquired priors (i.e., instructions), and (c) visual cues, are combined to generate the percept. We compared different Bayesian and circular inference (CI) models for their ability to fit the data. We particularly sought to understand whether circularity and aberrant correlations between priors and sensory evidence significantly contribute to the way we perceive the world.

## Results

To determine the effects of prior knowledge and sensory evidence in an ambiguous perceptual context, 55 participants were exposed to continuous presentation of a NC. The dominant percept was discontinuously sampled according to the procedure presented in **Figure 2** and was analyzed in terms of *relative predominance* (RP). RP corresponds to the overall probability of perceiving the SFA or SFB interpretation. A value of 1 or 0 would correspond to the SFA or SFB interpretation, respectively, fully dominating perception. A value of 0.5 would characterize a purely chance level wherein the 2 percepts are equiprobable.

**Figure 2:**
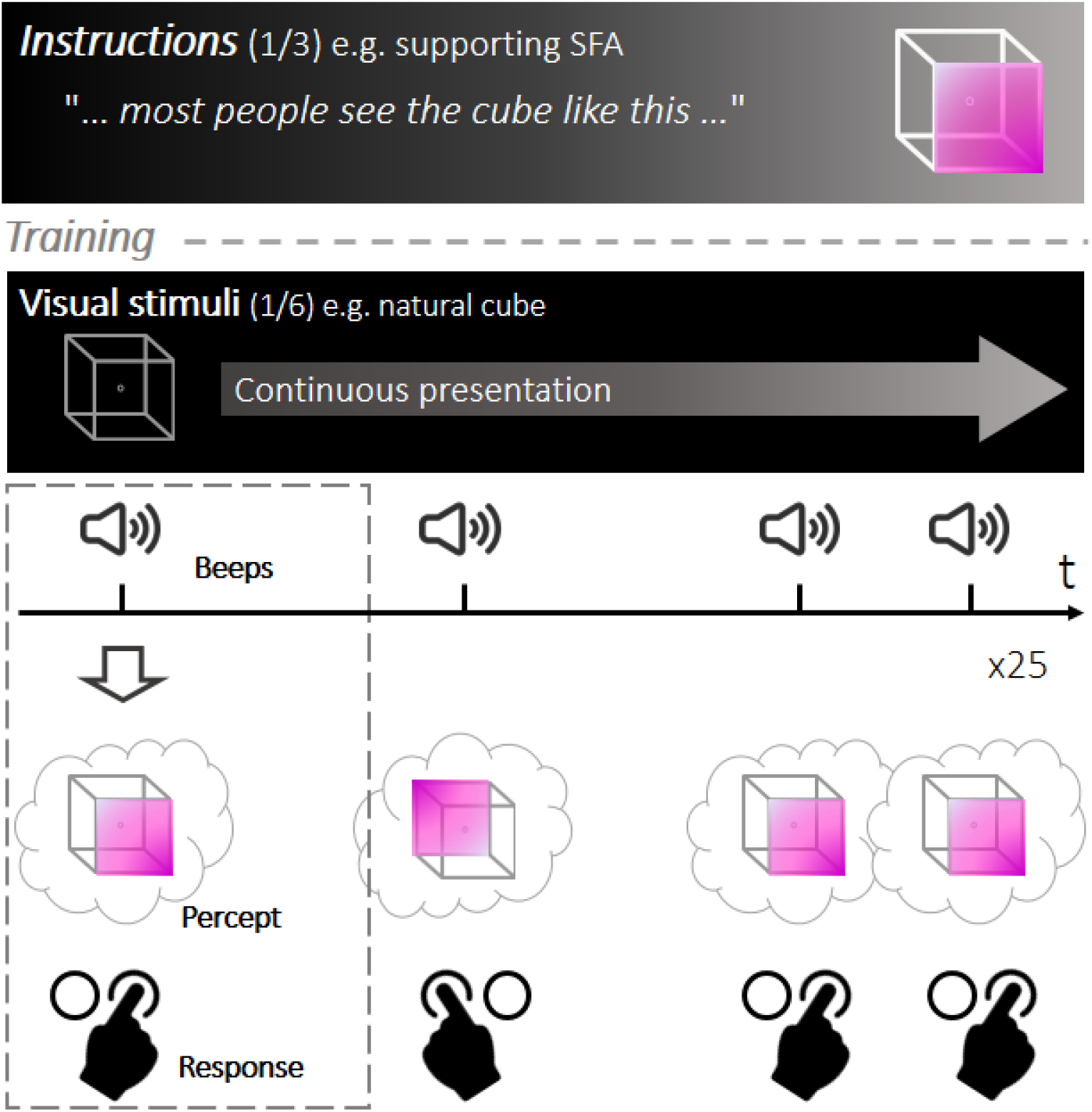
Experimental design. The task was inspired by [29]. Instructions were given at the beginning of the experiment (each participant received one set of instructions, creating a between-subjects design) and were followed by a short training phase to familiarize participants with the stimulus and the switches. During each run, one version of the cube was continuously presented to the participants, who were asked to discontinuously report their dominant percept by pressing a button every time a sound was heard. Each run consisted of 25 sound-trials (mean inter-sound-interval = 1.5 s). The main experiment consisted of 30 runs separated into 6 blocks of 5 runs each. In each block, a different variant of the stimulus was used. The first and fourth blocks always contained the ambiguous cube. The four cue conditions were randomly assigned to the four remaining blocks.

Sensory evidence was manipulated by casting the cube into shadow in such a way that it either contradicted or supported the SFA implicit prior (see stimuli, **Figure 1a (2-3,5-6)**). Visual cues were either strong or weak so that the analysis could reconstitute a cue pseudo-continuum from *strongly contradicting* to *strongly supporting.* In the completely ambiguous condition, no difference existed in the color of the two sides of the cube (see stimuli, **Figure 1a (1)**).

Prior knowledge was manipulated by randomly allocating participants to 4 groups. The first group was exposed to a tilted cube, which was expected to neutralize the SFA implicit bias (**Figure 1a (4)**). The remaining 3 groups viewed a normal cube but received different explicit instructions that either “supported”, “contradicted”, or were “neutral” with respect to the SFA bias (**Figure 1 b**).

### Model-free analysis

The effects of prior knowledge and sensory evidence manipulation are presented in **Figure 3**. RP was not significantly different between the 2 ambiguous blocks (runs 1-5 and 16-20) in any of the groups (**p > 0.1**), indicating only minor effects of fatigue (at least until the 20th run) and a stable effect of the instructions. Manipulation of sensory evidence significantly impacted bistability, with RP increasing as the visual cue changed from strongly contradicting to strongly supporting (**β = 0.415, p < 0.001**). Manipulation of prior knowledge through instructions only affected RP in the case of contradicting instructions, with a significant overall reduction in RP (**β = −0.096, p < 0.001**). Tilting the cube in the absence of any instruction resulted in a significant decrease in RP (**β = 0.103, p < 0.001**), which substantiated the effect of an implicit prior that naturally biases perception toward SFA dominance. Importantly, we found a significant interaction between the continuous effect of cue and the effect of contradicting instructions (compared to the normal cube with supporting instructions and the tilted cube with no instructions; **β = 0.265, p = 0.016** and **β = 0.265, p = 0.021**, respectively). Note that this interaction should not be present for a purely Bayesian observer, since the contribution of sensory evidence and priors (when expressed as the log odds ratio) should be additive.

**Figure 3:**
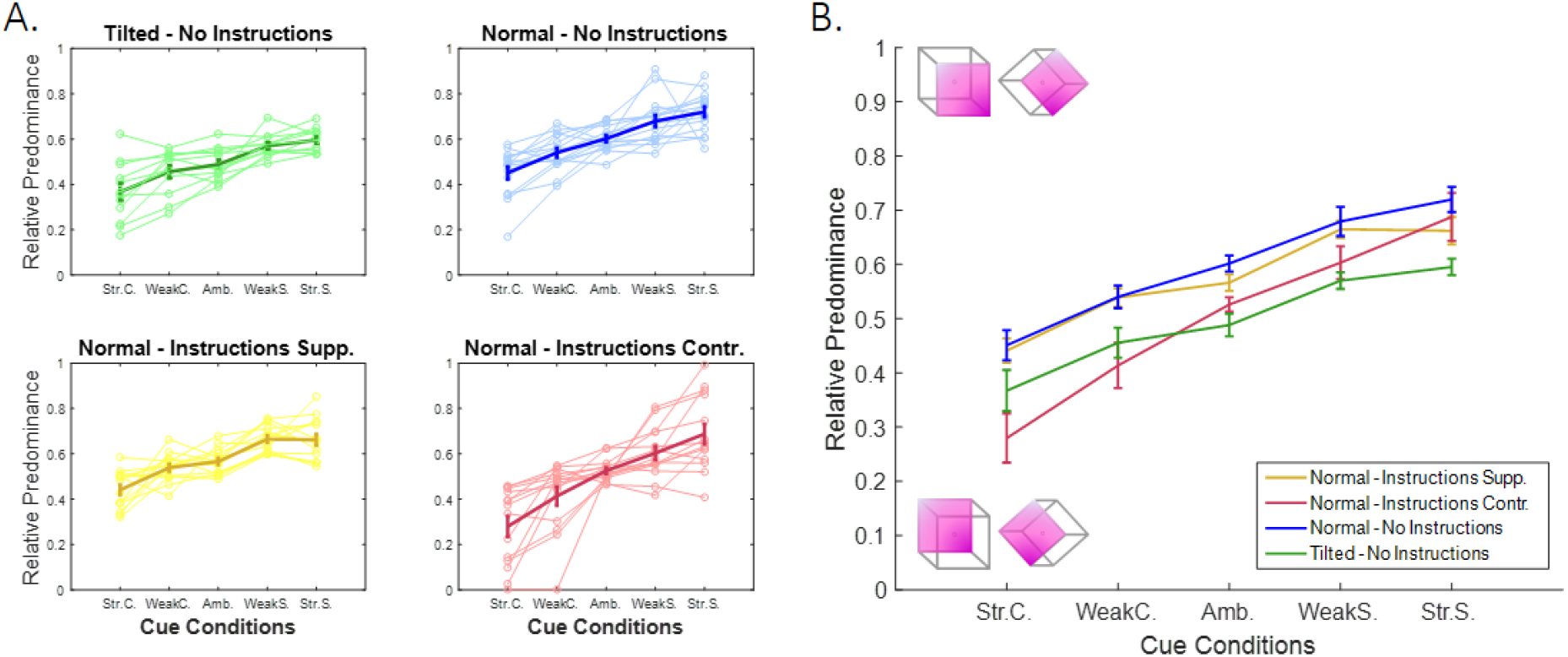
Relative predominance between conditions. **(a)** The four subplots illustrate the four different prior conditions: tilted cube (top left, green; N=12) or normal cube with no instruction (top right, blue; N=15), supporting instructions (bottom left, yellow; N=14) or contradictory instructions (bottom right, red; N=14). The x-axis presents the 5 cue conditions, ranging from strong cue supporting the SFB interpretation (left) to strong cue supporting the SFA interpretation (right). Thin lines correspond to the behavior of single participants (outliers are not presented), and thick lines represent the average RP for each group, after removing the outliers (±SE). **(b)** Between-groups comparison of average RP. A linear mixed-effects model revealed significant effects of sensory evidence (**p < 0.001**) as well as the prior (contradictory instructions, **p < 0.001**) and tilt (**p < 0.001**)) manipulations. We also observed a cue x instruction interaction for the contradictory instructions (red curve) compared to supporting instructions (yellow curve, **p = 0.016**) and the tilted cube (green curve, **p = 0.021**). See also **Figures S6, S7**.

### Model-based analysis

To test our hypothesis that circularity shapes bistable perception, we fitted a CI model to the average data, similar to the one introduced by Jardri and colleagues [26]. This model assumes that participants perform approximate inference due to the reverberation of sensory evidence and priors in the hierarchy as a result of unbalanced inhibitory control (**Figure 4a, right panel**). Furthermore, we compared the performance of our CI model against that of 2 Bayesian models performing exact inference: first, a naïve Bayes (NB) model, which is identical to the multiplicative rule of Bayes’ theorem (**Figure 4a, left panel**), and second, a weighted Bayes (WB) model in which different levels of trust (weights) could be assigned to sensory cues and priors. The WB model was equivalent to a NB model in which all the weights were set to 1 and equivalent to a CI model without any reverberating messages (**Figure 4a, middle panel**). The NB, WB and CI models can thus be considered 3 versions of the same model with an increasing number of parameters being fitted to the data. Predictions for the 3 models are presented in **Figures 4b and 4c**.

**Figure 4:**
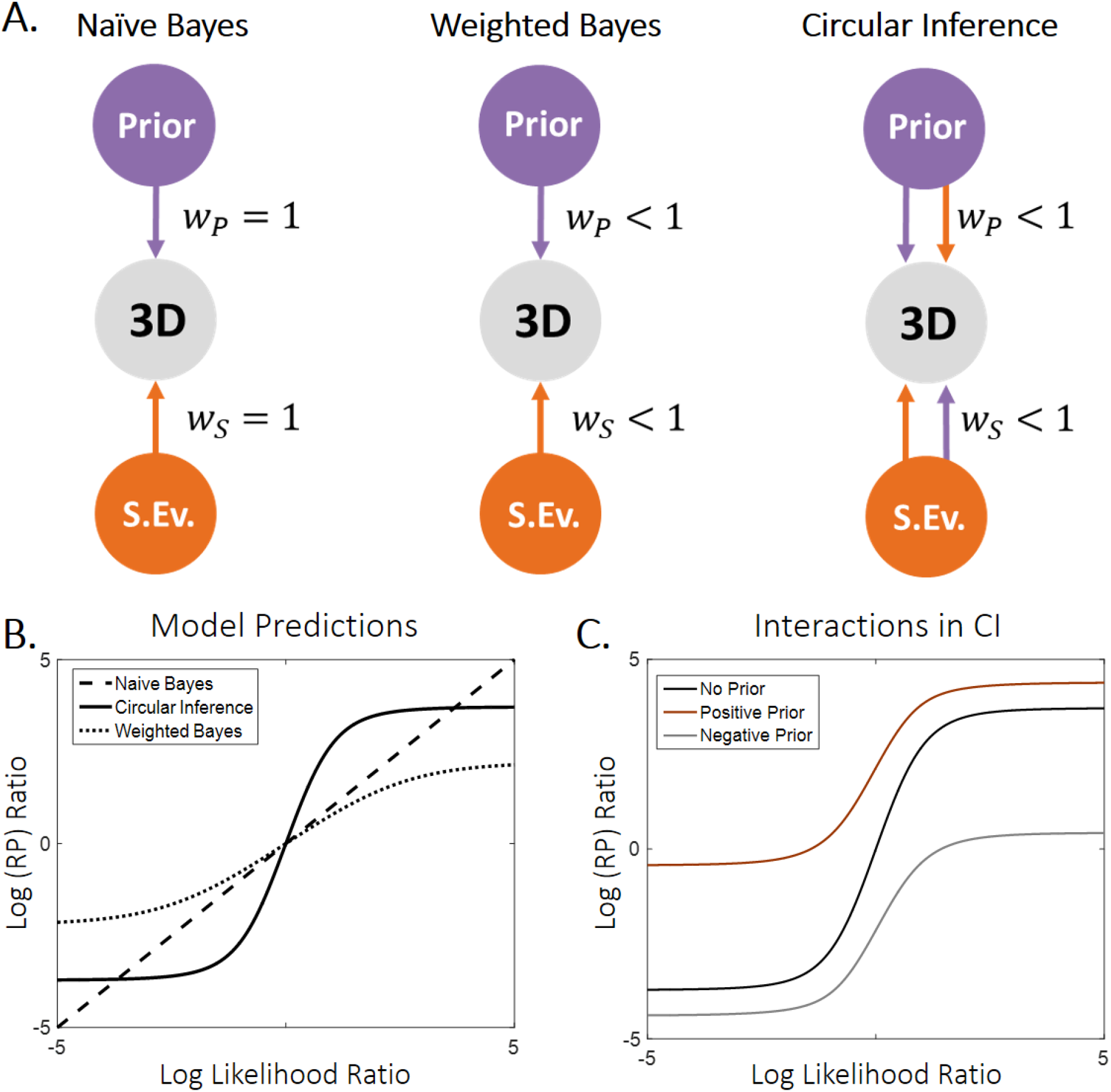
Illustration of models and model predictions. **(a)** Three different models were used to fit the data. The simplest model (naïve Bayes (NB), **left panel**) consisted of a simple addition of the sensory evidence and prior on the log scale and is equivalent to a three-layer generative model in which all the connections are equal to 1. The weighted Bayes (WB) model (**middle panel**) further assumes that there is only partial trust between the nodes of the generative model. Importantly, both the NB and WB models perform exact inference. Finally, we used a circular inference (CI) model (**right panel**) that further allows reverberation and overcounting of sensory evidence and prior knowledge. (b) Log(RP) ratio predicted by the models as a function of the log-likelihood ratio. The NB model predicts a linear dependence, whereas both the WB and CI models predict sigmoid curves (due to the saturation imposed by the weights). Furthermore, the 3 models make different predictions about the slope of the curves around zero. The NB and WB models predict a slope of 1 and less than 1, respectively, and only the CI model predicts a slope greater than 1. (c) In the CI model, the slope of the log-likelihood/log-posterior curve also depends on the log-prior as a result of the reverberations, indicating an interaction between the two different types of information [27]. Weaker priors are associated with steeper sigmoid curves. See also Figures S1 – S5.

**Figure 5** illustrates the best-fitting NB **(5a)**, WB **(5b)** and CI models **(5c). Figure 6** presents the values of the free parameters in the 3 models. The 3 models predict very different values for likelihoods and priors. These differences can be easily explained by the NB model assuming perfect trust in sensory evidence and priors, whereas the other 2 models predict much lower weights (*w_s_* = 0.77, *w_P_* = 0.59 for the WB model and *w_s_* = 0.66, *w_P_* = 0.59 for the CI model).

The NB model captures most trends in the data qualitatively, with the following exceptions. First, it underestimates RP in the case of the normal cube without instructions (**Figure 5a, blue curve**), and second, it is unable to predict the correct slopes. The latter limitation is especially striking in the case of a normal cube with contradicting instructions, where the slope is larger than predicted (i.e., larger than 1; **Figure 5a, red curve**). The WB model performs better than the NB model in most conditions, but it also underestimates the effect of the cue when the instruction contradicts the SFA preference (see **Figure 5b, red curve**). In contrast, the CI model captures this last trend (see **Figure 5c**), suggesting that the variability of the cue effect (the slope) in different conditions is due to circularity in the inference process.

**Figure 5:**
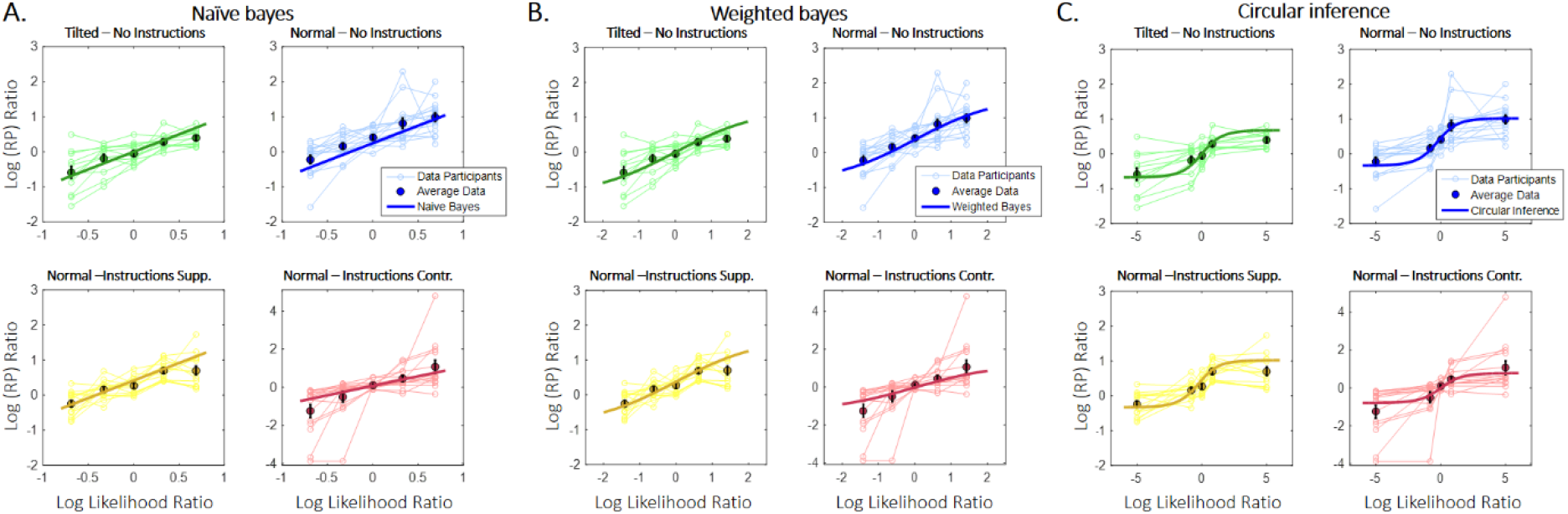
Observed and predicted log(RP) ratios as a function of the log-likelihood ratio. Different colors correspond to different prior conditions. Thin lines represent single participants’ data, highlighted points correspond to average RP (±SE), and thick lines illustrate model predictions. The three models are presented separately, since likelihood was itself considered a free parameter [**(a)**: NB, **(b)**: WB, **(c)**: CI]. The models were fitted to aggregated data from all participants by minimizing the mean squared distance between the observed and predicted log(RP) ratios. See also **Figures S1 – S4**.

**Figure 6:**
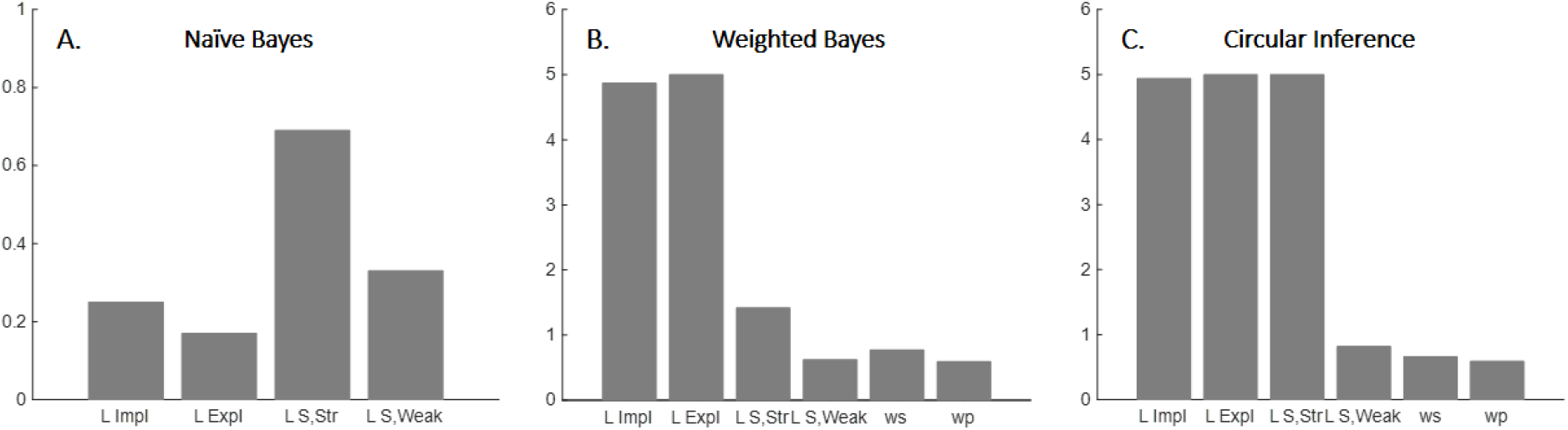
Optimal values of free parameters for the three models [(a): NB, (b): WB, (c): CI]. The NB model had fewer free parameters than the other 2 models, since the two weights were by definition fixed to 1. We observed important differences in the values of the likelihoods (L_S,Str_, L_S,Weak_) as well as in the values of the priors (L_impl_, L_expl_) between the NB model, on one hand, and the WB and CI models, on the other hand. These differences were mainly due to different values for the weights (w_s_, w_P_). See also **Figures S1 – S4**.

A quantitative comparison of the 3 models using BIC scores, which penalizes the use of extra free parameters in the WB and CI models, indicated that the CI model significantly outperformed the 2 Bayesian models (BIC scores for NB = −242.65, for WB = −240.77, and for CI = −249.49). A lower BIC score indicates that the model better fits the data, with a difference larger than 2 considered positive and a difference larger than 6 considered strong *(δ_BIC_* = 6.84 for comparison of the CI and NB models and *δ_BIC_* = 8.72 for comparison of the CI and WB models).

## Discussion

The goal of the current study was to decipher how priors and sensory evidence are combined to shape bistable perception. We particularly wished to investigate whether such integration is probabilistically optimal or if other principles are at play, contributing to the debate on whether bistable perception is a by-product of perceptual inference (regardless of its neural implementation). Our results suggest an imperfect neural implementation of probabilistic inference, possibly due to an imbalance between excitation and inhibition in neural circuits.

As previously reported, we found an asymmetry in the way participants interpreted the completely ambiguous NC [8]. This finding supports the notion of an implicit preference (implicit prior) to perceive objects in an SFA configuration [3]. More surprisingly, we showed that this preference could be explicitly manipulated by giving information that either confirmed or rejected it (explicit prior). In agreement with previous studies [28–30], adding visual cues also significantly biased perception toward the corresponding interpretation. The qualitative effects of implicit priors, explicit priors and sensory evidence appeared compatible with a probabilistic combination of information, suggesting that Bayesian inference was still at work.

However, we also found a significant interaction between priors and sensory evidence that could not be explained by exact inference. In particular, the effect of sensory cues was stronger when the prior was more ambiguous (e.g., when the implicit preference for SFA was contradicted by instructions) and weaker in the absence of a prior (e.g., a tilted cube). In contrast, Bayes’ theorem predicts that sensory cues are weighted according to their reliability, independently of the prior. Through parametric model comparison, we found that the present data could be better accounted for by a CI model, in which prior beliefs (i.e., the instructions and SFA preferences) corrupt new sensory evidence (i.e., ambiguous cues are misinterpreted as supporting the current belief) and *vice versa.* This corruption could be the result of feedback to sensory areas insufficiently controlled by inhibition [22]. Such feedback could also cause multistable perception (i.e. generate a bistable attractor; see **Supplementary Figure S5**) by temporarily stabilizing the current percept despite the absence of supporting evidence [27].

These findings add new elements to a long-lasting debate in neuroscience that questions whether perception is mostly driven by bottom-up processes, or whether top-down effects are equally important [21]. Multiple studies have investigated how low-or high-level manipulations affect bistability, without offering definitive answers. For the former, authors have used priming or suppressing effects (usually attributed to adaptation) [31–34], changes in retinal location [35], manipulation of the type of presentation (continuous-intermittent) [36, 37], and direct manipulation of the properties of the stimulus, like intensity [38] and completeness [39]. In contrast, studies of high-level manipulations have focused on the effects of volition [40, 41], expectation and prediction [42], attention [43–45], learning [46], mental imagery [47], knowledge of reversibility [48] and finally the preference for stimuli with a statistical structure similar to that of natural images [8,49,50]. Note however that the present study was not designed to test specific neural mechanisms such as adaptation and noise.

Consistent with the present study, some authors have focused on how these various effects are combined [51–53]. Moreno-Bote et al showed that cue combination in a bistable display can be well explained by a multiplicative law (their predictions are similar to the NB model described here) [54], whereas Zhang and colleagues demonstrated that different types of priors are effectively combined [4]. Here, we have gone a step further and investigated how top-down (prior manipulation) and bottom-up (sensory cues) effects interact. Rather than inducing a prior through learning, as is widely done in the literature [46, 47], we directly manipulated participants’ expectations. This manipulation assumes that instructions can generate a high-level prior affecting perceptual processing in a way similar to a learned prior (as in [55])

Despite the amount of available data and the apparent simplicity of the problem, very few studies in the literature have applied normative explanations for bistable perception that include data-fitting [54]. Although proposing a complete model of bistable perception based on circular inference goes beyond the scope of the paper, our present results suggest that a local message passing algorithm with the addition of information loops could constitute the basic principle of such a normative model. Some alternative normative models have relied on a simplified form of Markov Monte-Carlo sampling. More precisely, they assumed that the current percept is based on taking one sample from the posterior distribution and using this sample as a prior for the next time step [56, 57]. However, Markov Monte-Carlo sampling requires very long sampling times (because of temporal correlation between samples) to perform accurate inference. A possible argument in favor of circular inference would be that it can reach correct conclusions quickly and accurately in most perceptual tasks, except in particularly ambiguous cases [22], making circular inference a powerful model for perceptual inference in unambiguous cases.

From a methodological point of view, and in contrast to most studies on bistable perception, in which participants continuously report the dominant percept with a sustained button-press [58, 59], we asked participants to respond discontinuously, after being exposed to a go-signal [29]. This procedure has two main advantages. First, it minimizes the role of attention. Indeed, it has been shown that attention plays a crucial role in bistable perception, especially for certain bistable stimuli [41, 60]. The inability to control for differences in attentional load between participants could be an important source of uncertainty and even partly explain the huge variability usually observed in some publications (see [29]). Second, this procedure is less affected by differences in reaction time, as one could use the time of the sound as a proxy for the time of the decision. As a consequence, discrete sampling not only seems ideal for a rigorous experimental exploration of bistable perception but is also useful for adapting this task to specific clinical populations with well-known attentional and motor problems.

Finally, some limitations need to be acknowledged. First, because of the type of priors used (instructions), we were obligated to use a between-subjects design, which prevented us from comparing the effects of different instructions in the same participant. As a result, there were only 5 conditions per participant, and we could only fit our models to averaged data, ignoring variability between participants (see also [15, 54]). Second, all the models under consideration were based on an assumption of temporal independence between the percepts at the time of the sounds. This assumption can be partly justified by the weak autocorrelation of the averaged data (see **Supplementary Figure S6**), although these autocorrelations may be stronger in individual participants [56]. Nevertheless, temporal statistics would not affect the qualitative predictions of the models [54]. In particular, temporal statistics without circular inference would not provide a valid alternative to the present findings, including the slopes and the cue × instruction interaction. Third, a response bias could partially account for instruction effects (explicit priors). However, a response bias would have a similar effect over responses across different cue conditions while leaving perceptual processing completely unaffected. Although the above is a possible interpretation of the data, it remains highly improbable, given the non-linear interaction found between instructions and visual cues (see also **Supplementary Figure S7** for additional arguments).

Overall, this study confirms that circular inference can be observed to a certain degree in healthy individuals. This unprecedented observation opens a range of crucial questions that suggest opportunities for further research: in what other ways could circularity affect cognition, and what are its neural substrates? Crucially, we must determine under what circumstances circular inference generates aberrant beliefs or percepts, such as those observed in pathological (neurological or psychiatric) contexts.

## STAR Methods

### Contact for reagent and resource sharing

Further information and requests for resources and reagents should be directed to and will be fulfilled by the Lead Contact, Renaud Jardri.

### Experimental model and subject details

Participants were healthy volunteers meeting the following inclusion criteria: age > 18 years, provision of informed consent, normal or corrected-to-normal near visual acuity, no past or current medical history of neurological or psychiatric disorders, and no current or recent use of psychotropic medication or toxic drugs (the demographic characteristics can be found in **Table 1**). Near visual acuity was quantified using the Parinaud score; we considered values less than or equal to 2 to be normal. Of the 65 participants initially recruited, 10 were excluded because of outlying mean RP values (more details are provided in the **“Quantification and Statistical Analysis”** section). We highlight that 7 of the 10 excluded participants also exhibited qualitatively bizarre behavior (such as opposite effects of visual cues), indicating a misunderstanding of the instructions or low attention levels. The study was approved by an ethics committee.

**Table 1:** Demographic characteristics of the 4 groups (without outliers). The 4 groups did not differ in terms of age, education or sex. ◊; F-test, ◦: Chi-squared test

| Variables | Tilted<br>(n = 12) | Instr. Supp.<br>(n = 14) | Instr. Contr.<br>(n = 14) | No Instr.<br>(n = 15) | Comparison |  |
| --- | --- | --- | --- | --- | --- | --- |
|  |  |  |  |  | Test | p |
| **Age**<br>mean (sd) | 23.33<br>(2.77) | 28.64<br>(7.19) | 28.93<br>(9.60) | 29.27<br>(11.73) | 1.31♦ | 0.28 |
| **Education**<br>mean (sd) | 17.25<br>(2.42) | 19.07<br>(1.94) | 18.57<br>(2.17) | 18.00<br>(1.96) | 1.77♦ | 0.16 |
| **Sex ratio** (m:f) | 3:9 | 7:7 | 8:6 | 9:6 | 3.87° | 0.28 |

### Method details

The general procedure (**Figure 2**) was inspired by Mamassian and Goutcher’s protocol [29] and consisted of 6 blocks of 5 consecutive runs. During each run, a 200 x 200 pixel NC displayed in the middle of a black screen was continuously presented to the participants. Using a forced-choice method, we asked participants to report their ongoing interpretation as soon as they heard a warning sound, which occurred 25 times in a pseudo-regular manner (mean inter-sound interval = 1.5 s, uniformly distributed between 1 and 2 s). Each response corresponded to a trial, providing a discontinuous sampling of the task’s perceptual dynamics. Runs were separated from one another by a black screen with a duration of 10 s to minimize between-run influences. The experiment was also interspersed with 5 between-block breaks of non-predefined duration. Prior to the experiment, participants were informed that they would be presented with empty cubes, the 2 possible interpretations of which were explicitly mentioned. The basic instruction was to passively view these cubes without trying to constrain perception.

We manipulated sensory evidence either by making the cubes homogeneously gray (i.e., perfectly ambiguous) or cuing them by shadows (**Figure 1a (1-3,5-6)**). This additional depth information was intended to bias perception toward one interpretation or the other. It was specified by two parameters. First, its orientation was defined in relation to the implicit prior. A shadow falling on the top left corner was expected to emphasize the SFA preference and thus was classified as a supporting cue. Conversely, a shadow that fell on the bottom right corner was characterized as a contradictory cue, as it went against implicit bias. Second, the strength of the cue (which can also be conceived in terms of the amount of sensory information) was controlled by the shadowing contrast level. Weak and strong cues corresponded to 20% and 30% contrast, respectively. The 1^st^ and 4^th^ blocks always consisted of presentation of an ambiguous cube. The other blocks were randomly allocated a different type of cue, defined by the 2 x 2 factorial combination of 2 possible orientations (contradicting or supporting) and 2 possible strengths (weak or strong).

Participants were separated into 4 groups (n = 12, 14, 14, and 15) that differed in terms of how we altered their prior knowledge. The first group was presented with a tilted cube, which was expected to neutralize the SFA implicit bias (**Figure 1a (4)**). The remaining 3 groups viewed a normal cube— where the implicit prior is deemed present—but received different types of instructions, which we used to manipulate their implicit prior. In Group 2, instructions explicitly mentioned the presence of the implicit bias:

> *“When looking at the cube, most people tend to see it with its front side on the right. Differently said, there is a natural tendency to see the cube mostly “from above”. In the present experiment, we aim to study the characteristics of this spontaneous preference.”*

Because the statement was correct, the instructions were considered to support the spontaneous bias (supporting instructions). In Group 3, participants were informed about a natural tendency to see the cube primarily as though it were seen from below. The wording was similar, but the statement was incorrect, thus contradicting the implicit prior (contradictory instructions). In Group 4, the participants received no complementary information. In this case, their prior knowledge could be considered akin to the implicit bias (neutral instructions). Note that, to avoid any additional priming effects, the difference among the 4 groups was only verbal, while all groups received the same visual instructions, including the stimulus and the 2 possible interpretations. As shown in Table 1, the 4 groups did not significantly differ in terms of demographic characteristics.

To neutralize the potential confounding bias of eye-movements, participants were additionally instructed to gaze at a fixation point in the middle of the screen. A training session allowed each participant to familiarize himself/herself with the stimuli and the apparatus.

The experiments were implemented in MATLAB v. 2011b (MathWorks, Natich, MA), using Psychtoolbox v. 3.0.10. Stimuli were displayed on a 17-inch LED screen with a resolution of 1280 x 1024 pixels. Responses were collected using a classical computer keyboard. A chin-cup and forehead bar ensured immobilization of the participant’s head at a distance of 60 cm between the eye and the screen.

#### Model-free analysis

RP was calculated by taking the grand mean of responses across trials, runs and participants. It can be interpreted as the general probability to perceive one interpretation or the other on each trial.

#### Model-based analysis

We conceptualized perception as an inferential process, in which the brain generates a subjective belief about the possible interpretations of the NC (i.e., a posterior probability) and uses it to make a perceptual decision, particularly whether it is an SFA or SFB cube. Three different models were fitted to the average RPs of the 4 groups. All the models assumed independence between the sequential perceptual decisions within a run. They differed in how the 3 main effects of the experiment (sensory evidence *S,* an implicit prior *P_impl_*, and an explicit prior *P_expl_*) were combined to give rise to the posterior probability *P*(*X|S,P_impl_,P_expl_*). In this expression, *X* is a binary variable that corresponds to the 3D interpretation (*X* = 1 corresponds to SFA, *X* = 0 corresponds to SFB.

The simplest model that was fitted to the data is the NB model, which assumes perfect integration of likelihoods and priors according to the Bayes theorem. Consequently, it’s equivalent to a basic multiplicative rule [54] (additive rule in the log scale) (eq. 1; **Figure 4a, left panel**). The WB model extended the NB model by assuming only partial trust to the sensory evidence and prior information (eq. 2; **Figure 4a, middle panel**). Crucially, both models are Bayesian models performing exact inference. Finally, the third model is a *circular inference model* [26], meaning that information is not only weighted, as in the WB model, but it’s also amplified, due to information loops (eq. 3; **Figure 4a, right panel**). As a result, the CI model is doing sub-optimal inference, which renders it qualitatively different from the other 2 models.

The 3 models are quantitatively described by the following equations:

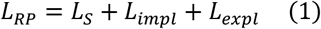

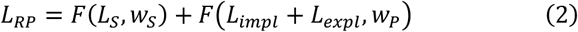

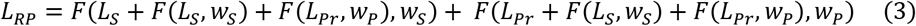

where *F(L,w)* is a sigmoid function:

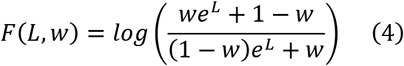

and *L_Pr_ = L_impl_ + L_expl_. L_RP_* corresponds to the log-ratio of the RP and is taken to be equal to the log-posterior ratio. That assumption is because we assume that perceptual decisions are made using probability matching, a commonly observed strategy in sequential 2AFC tasks [20,54,61]. We note that applying a SoftMax to the log posterior odds (a more appropriate model for perceptual decisions) would only induce a global change in the gain of the former and would not affect any of our conclusions.

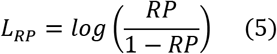

The log-likelihood ratio *L_s_*, the implicit log-prior ratio *L_impl_* and the explicit log-prior ratio *L_expi_* are given by the following equations:

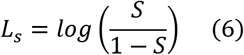

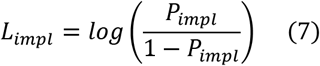

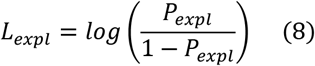

Because none of these variables was known, they were all treated as free parameters. To reduce as much as possible the total number of free parameters that needed to be optimized, we further considered symmetry both in the effects of the cues and the instructions, resulting in 4 free parameters Finally, *L_s,trong_, L_s,weak_,L_impl_,L_expl_*).

Finall, *w_S_* and *w_P_* (appearing only in the WB and CI models) correspond to participants’ trust (or weight) in the sensory evidence and priors, respectively, and constituted the 2 additional free parameters of those models:

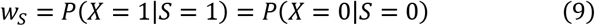

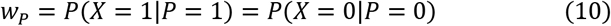

Importantly, since the SFA prior was completely uninformative in the case of the titled cube, we considered the following:

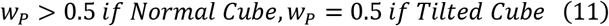

As a control, we also considered the case in which *w_P_* has the same value in all conditions (see **Supplementary Figure 1**).

An illustration of the different models is presented in **Figure 4a** The CI model (**Figure 4a, right panel**) hypothesizes that the perceptual system performs approximate inference due to unbalanced inhibitory control. Those impairments lead to a failure to remove efficiently redundant messages: a reverberating prior, which is misinterpreted as sensory evidence, re-ascends the hierarchy and corrupts the likelihood term and redundant sensory evidence, which descends the hierarchy and corrupts the prior term. Additionally, as in [26], a cross-term is added to each component, rendering likelihood and prior information completely inseparable. Because of those extra terms, the sensory evidence and prior components become aberrantly correlated, and consequently they generate an interaction (**Figure 3c**; [27]). Note that the WB model (**Figure 4a, middle panel**) can be derived from the CI model by removing the reverberated terms, while the NB model (**Figure 4a, left panel**) by further assuming: *w_s_ = w_P_* = 1.

The CI model used here was similar to the model used by Jardri and colleagues to explain participants’ behavior (both those suffering from schizophrenia and healthy participants) in a probabilistic reasoning task [26]. Nevertheless, an important difference needs to be highlighted. In the present study, the redundant messages corrupted the original messages only once (there was still overcounting of information, but the amount of amplification stayed constrained), which is equivalent to setting *a_s_* and *a_P_* (the parameters in the original model that represented the number of times the redundant messages were taken into account) equal to 1. The reason was twofold. First, fixing the number of loops did not change the results qualitatively. Indeed, the resulting model predicted both a slope larger than 1 and an interaction between sensory evidence and priors, the two characteristic features of circular inference observed in the data. Second, the additional complexity (2 more free parameters) did not further improve the fit (see **Supplementary Figure S2**).

**Figure 4b** illustrates the predictions of the 3 models. Contrary to the linear NB model, both the WB model and the CI model are non-linear models, due to the saturation of the posterior that is caused by the weights. Importantly, the 3 models make different predictions about the slope of the log-likelihood/log-posterior curve around 0: the NB model and WB model predict a slope equal to and smaller than one, respectively. Interestingly, only the CI model can generate a slope that is larger than one, due to overcounting of the prior and of sensory evidence. Moreover, it predicts interaction between the prior and sensory evidence, such that the slope differs depending on prior strength and weight (**Figure 4c**).

Finally, in eq. 1–3, we assumed that instructions act as an additional prior term, essentially changing the strength of the implicit preference independently of the presence of a visual cue. As a result, any interaction between the effect of the cue and the effect of the instructions is forbidden under Bayesian formalisms and can only be explained by non-Bayesian mechanisms such as the presence of circular inference. It is worth noting though that alternative interpretations of the instructions (which are even more complex) might also generate such an interaction, notably likelihood-dependent instructions, or instructions that directly affect the reliability of the sensory evidence. Those additional models were also considered and compared to the CI model (see **Supplementary Figures S3 and S4**).

### Quantification and statistical analysis

#### Model-free analysis

Because RP is a ranged variable, we performed exclusively non-parametric analyses. The effects of priors, sensory evidence, and their interaction were tested using a linear mixed-effects model comprising the effects of cues and instructions as well as their interaction as fixed effects, together with Gaussian random effects for intercepts and slopes. For significant omnibus effects, we performed post hoc comparisons using either paired or unpaired rank-sum tests to clarify simple effects in the 2 x 2 design. Finally, one-sample Wilcoxon signed rank tests were performed to compare the mean RP with 0. 5, i.e., chance level. All participants whose RP was more than 1.5 interquartile ranges below the first quartile or above the third quartile were considered as outliers and were excluded. All significance tests were performed on the final sample of the 55 participants (12, 14, 14 and 15 for each group respectively), they were two-tailed and used an alpha value of 0.05 in the statistical toolbox of Matlab v. 2011b (MathWorks, Natich, MA). More information about the different tests can be found in the **Results** and in the figure legends.

#### Model fitting

All the models were fitted to the data by minimizing the mean squared distance between the log(RP) ratio for the different conditions and the predictions of the models. Instead of simply considering the means, we used data points from each participant, making full use of the available information but assuming that the parameters did not vary between participants. The optimal values for parameters were obtained using a non-linear programming method (sequential quadratic programming; a built-in MATLAB function), appropriate for non-linear constrained multivariable functions. To avoid local minima, the optimization process was repeated 100 times for each model, with initial values chosen each time randomly from the parameter space.

#### Model comparison

We compared the quality of the fits for the 3 models using BIC scores. We approximated the likelihoods of all the models as normally distributed. The BIC score can then be calculated by the following equation:

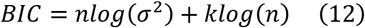

where *n* is the total number of data points (5 points per participant), *σ*^2^ is the mean squared error, and *k* is the number of free parameters (4 for NB, 6 for the other models).

## Author Contributions

Conceptualization: P.L., C-E.N., R.J., S.D.

Methodology: P.L., C-E.N., R.J., S.D.

Software: P.L., C-E.N.

Formal Analysis: P.L., C-E.N.

Investigation: P.L., C-E.N., M.E.

Resources: P.L., C-E.N., M.E., R.J., S.D.

Writing – Original Draft: P.L., C-E.N., M.E., R.J., S.D.

Writing – Review & Editing: P.L., C-E.N., M.E., R.J., S.D.

Visualization: P.L., C-E.N., R.J., S.D.

Supervision: R.J., S.D.

Project Administration: P.L., C-E.N., R.J., S.D.

Funding Acquisition: R.J., S.D.

## Acknowledgements

P.L. was supported by a PSL University PhD fellowship. S.D. was supported by an ERC consolidator grant “Predispike” and by the James McDonnell Foundation award ‘‘Human Cognition”. This research was also supported by: ANR-17-EURE-0017 FrontCog and ANR-10-IDEX-0001-02 PSL grants *(Département d’ Etudes Cognitives* of the *Ecole Normale Supérieure).*

