## Supplemental Information for "Circular inference in bistable perception"

### Supplementary Information

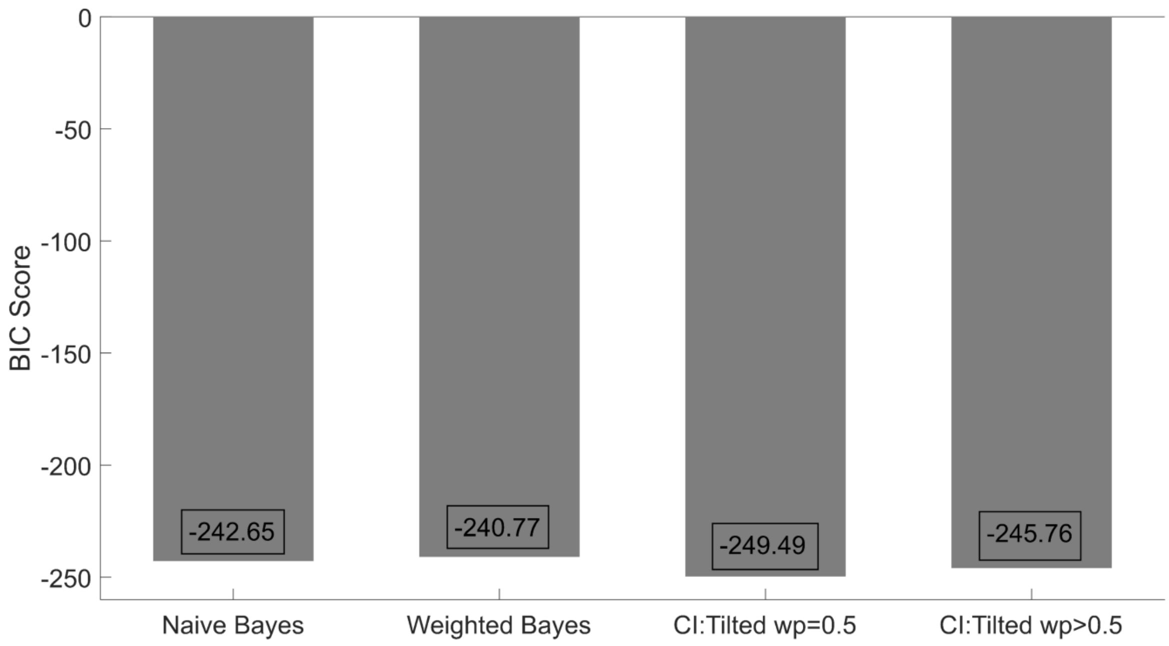


**Figure S1 (related to Figures 4, 5 and 6): Comparison of models assuming different effects of priors in the case of the tilted cube**. In the **main text**, we made the assumption that $w_P = 0.5$ when the cube is tilted, since in that case the prior becomes uninformative and thus irrelevant. An alternative would be that the prior weight remains larger than 0.5 (as in the case of the normal cube), and that only the log-prior ratio becomes equal to zero. The 2 alternatives cannot be differentiated in the cases of the NB and WB models, but they make different predictions in the case of the CI model, in which a reverberating likelihood term appears inside the prior term. This reverberating term disappears completely if we assume $w_P = 0.5$, whereas it remains if we assume $L_P = 0$. Formal comparison of the two models using the BIC score revealed that the former ($w_P = 0.5$) outperformed the latter ($w_P > 0.5$) (BIC scores: -249.49 vs. -245.76, respectively).

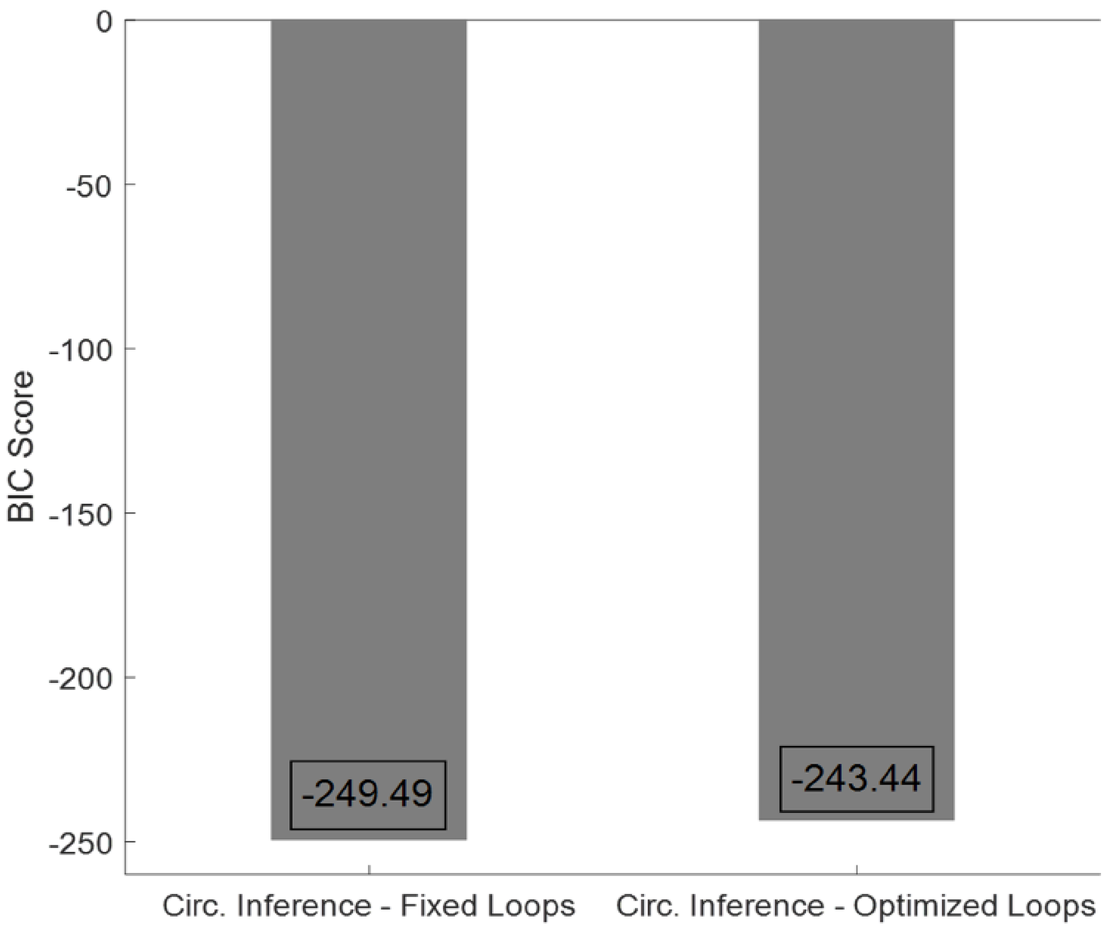


**Figure S2 (related to Figures 4, 5 and 6): Comparison of circular inference models with fixed and optimized loops**. In a previous study, Jardri and colleagues considered a *CI model* in which the strength of climbing loops (reverberation of sensory evidence, $a_S$) and the strength of descending loops (reverberation of priors, $a_P$) were considered free parameters and were optimized [1]. The predictions of the model were summarized by the following equation: $L_C = F(L_S + F(a_c L_S, w_S) + F(a_d L_P, w_P), w_S) + F(L_P + F(a_c L_S, w_S) + F(a_d L_P, w_P), w_P)$. In the current study, we fixed the values of these 2 extra parameters to 1, obtaining equation (1) (**Main Text**). The two models make the same qualitative predictions, as they both contain reverberating terms that render likelihood and prior inseparable. We quantitatively compared the two models using their BIC scores. We found that the simplified model (fixed loop strength) performed better than the full model (optimized loop strength) (BIC scores: -249.49 for the former vs. -243.44 for the latter).

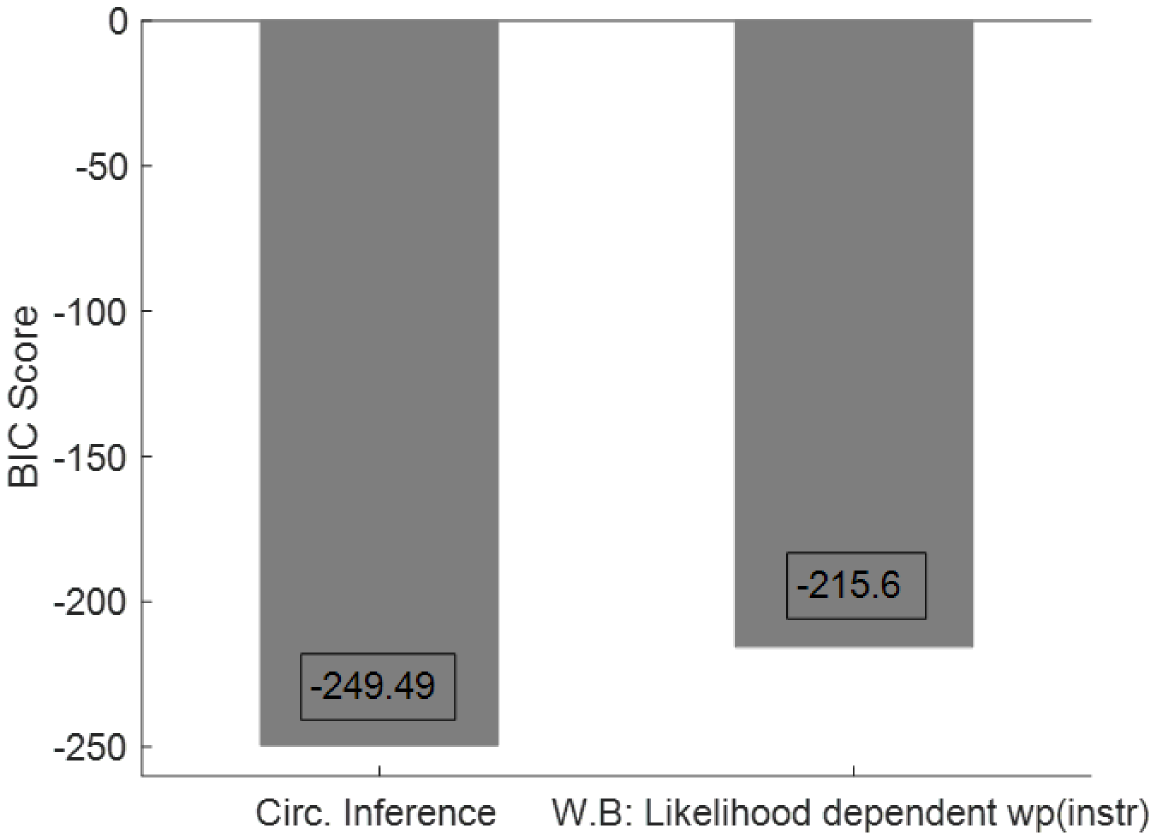


**Figure S3 (related to Figures 4, 5 and 6): Comparison of the CI model with a WB model in which the instructions are likelihood-dependent.** A likelihood-dependent effect of the instructions may constitute an alternative explanation for the interaction observed between the effect of the visual cues and that of the instructions/priors. In our framework, such an interpretation can be implemented as follows: $L_{RP} = F(L_S, w_S) + F\left(L_{impl}, w_{P,impl}\right) + F(L_{expl}, w_{P,expl}(L_S))$, where the weight attributed to the instruction is likelihood-dependent. Despite its plausibility, such an implementation drastically increases the complexity of the model, since it comprises 9 free parameters (instead of $w_P$ , we now have [$w_{P,impl}$, $w_{P,expl}\left(L_{S,amb}\right), w_{P,expl}\left(L_{S,Str}\right), w_{P,expl}(L_{S,weak})$]). A formal comparison of the CI model with this version of the WB model reveals a clear superiority of the former (BIC scores: -249.49 vs -215.6, respectively).

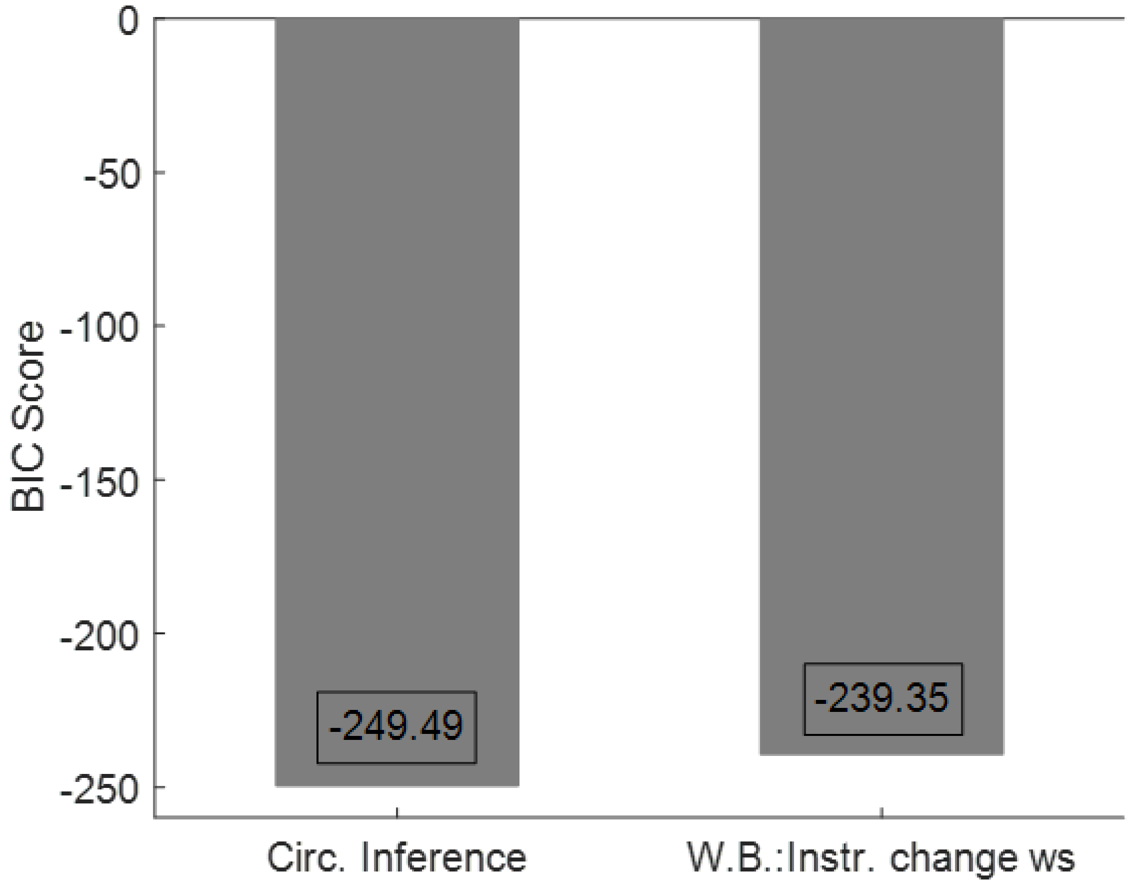


**Figure S4 (related to Figures 4, 5 and 6): Comparison of the CI model with a WB model in which the instructions directly affect the reliability of the visual cue.** The observed interaction between visual cues and priors could be explained by assuming that the instructions do not act as a prior but instead change the reliability of the sensory evidence. In our framework, such an interpretation could be implemented as follows: $L_{RP} = F(L_S, w_S(Instr)) + F\left(L_{impl}, w_P\right)$, where the sensory weight depends on the instructions. This model comprises 7 free parameters, and a formal comparison with the CI model reveals that circularity (plotted on the left) offers the best interpretation of the data (BIC scores: -249.49 vs -239.35, respectively).

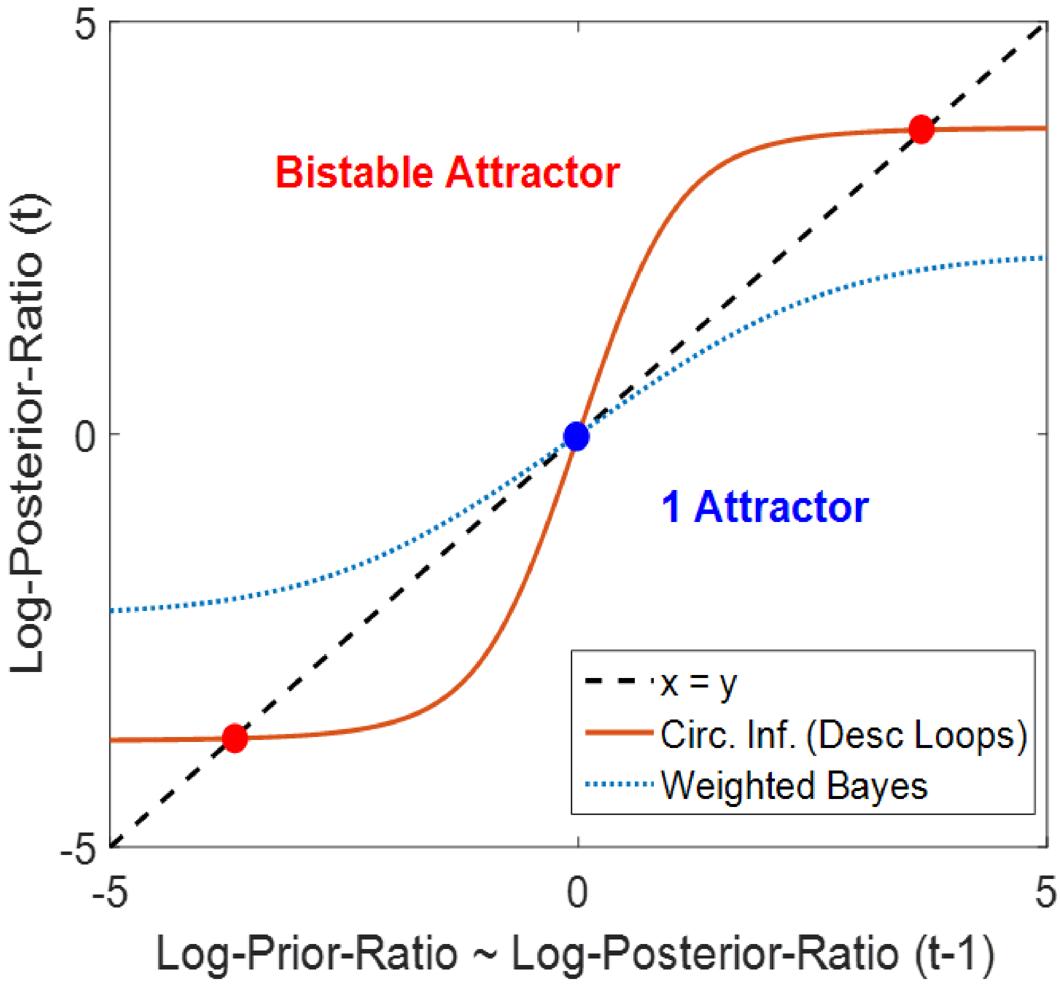


**Figure S5 (related to Figure 4): Descending loops can generate a bistable attractor.** The present results suggest that an aberrant correlation between sensory evidence and priors might be at play in bistable perception, shaping which interpretation we see and when. In the same context, it is important to highlight that a *Circular Inference* model, but not a purely Bayesian model (e.g., the WB model), seems compatible with the phenomenology of bistable perception. When taking into account the dynamics, descending loops (i.e., amplifying accumulated data) introduce a positive feedback to the system, which generates a bistable attractor (2 stable states consisting of strong beliefs, one for each interpretation; red solid line). In contrast, a Bayesian model corresponds to a leaky integrator, in which the belief is similar to chance (blue dotted line) [2] and appears unable to generate bistability.

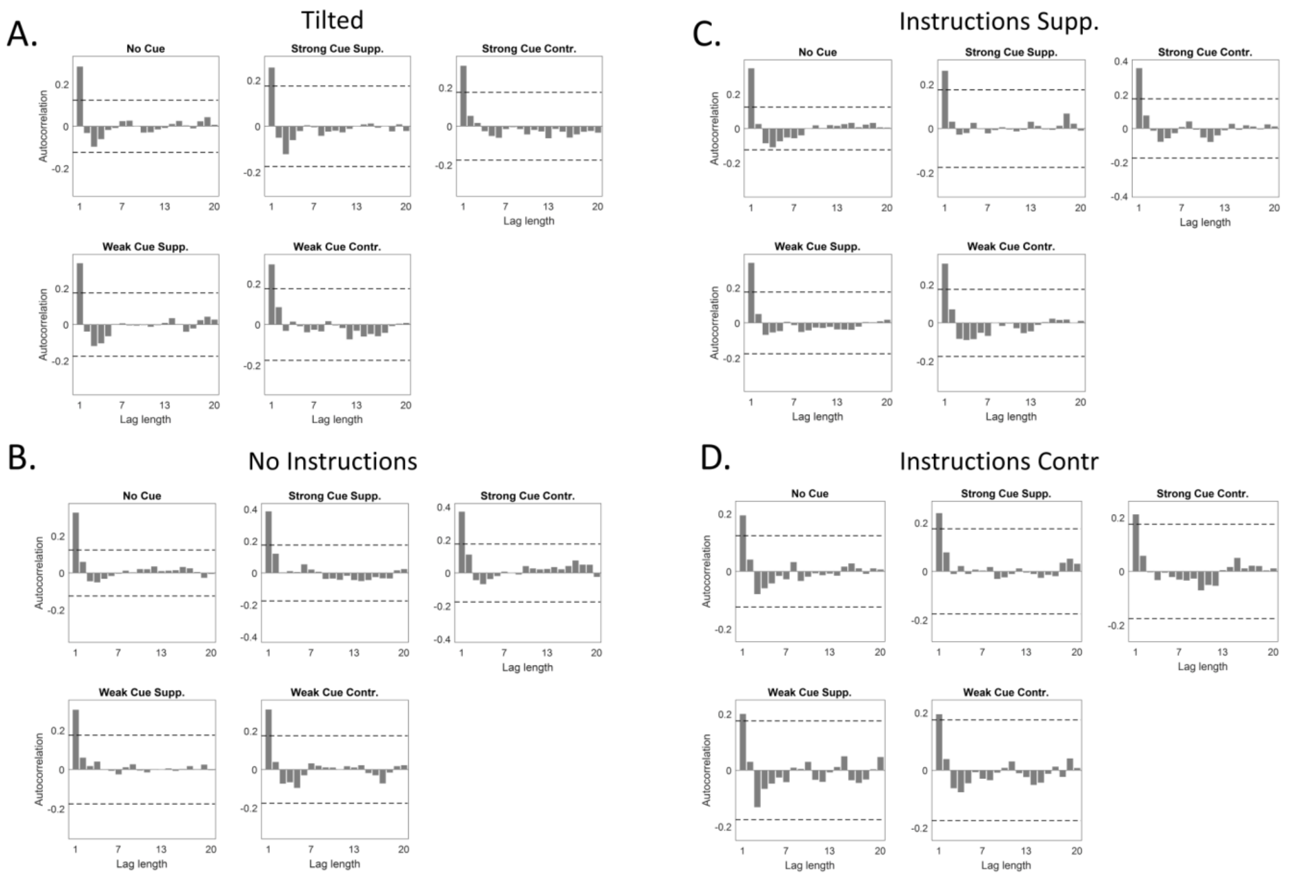


**Figure S6 (related to Figure 3): Average autocorrelation functions for the different cue conditions (different subplots) and the different prior conditions** [**(a)**: tilted group (N=12); **(b)**: normal cube, no instructions (N=15); **(c)**: normal cube, supporting instructions (N=14); **(d)**: normal cube, contradictory instructions (N=14)]. Dashed lines correspond to 95% confidence intervals for a white noise process. Interestingly, with the exception of small lags (lag = 1), no other point in any of the autocorrelation functions exceeds the dashed lines, meaning that the effects of history (at least for the average data) are restricted to small time differences and can be neglected. Consequently, none of our models took into account temporal statistics (see **Discussion** for further consideration of this issue).

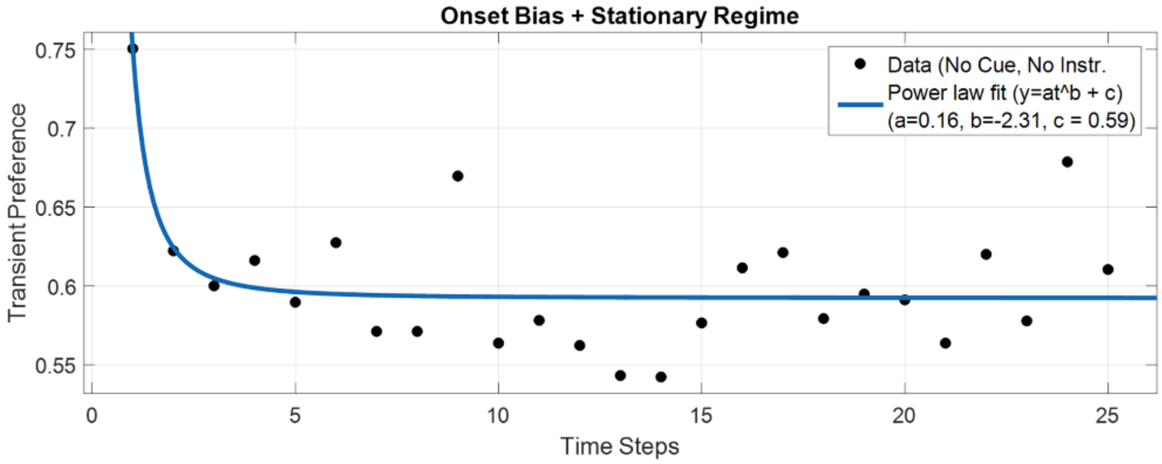


**Figure S7 (related to Figure 3): Transient preference (TP) as a function of time (black dots) and fitted power law model (blue curve)**. Transient preference can be defined as the RP for different time steps. As expected, TP attains a high value (TP(1) = 0.75) at the beginning of a run (t = 1), indicating the presence of an onset bias that rapidly decreases until reaching a stationary regime (TP(st. reg.) ~ 0.6) [3]. Interestingly, TP in the stationary regime is above chance, indicating the presence of a persistent bias even after the onset bias fades out (the implicit bias that we describe in the **Main Text** is a combination of those two biases). This figure is reassuring, since absence of this pattern would indicate either a response bias or a very long inter-stimulus interval. This figure corresponds to the normal cube/no instructions/no cue condition (N=15), but a similar pattern was obtained for all other conditions as well (not presented).
